# The level of workaholism and its relation to positive and negative perfectionism

**DOI:** 10.1101/604934

**Authors:** Basim Mohammad Ali Aldahadha

## Abstract

The present study aimed to test how common workaholism is and which groups are most targeted in the workplace among Jordanian employees. Additionally, the roles of positive and negative perfectionism in workaholism were investigated. The sample consisted of 686 employees. All of them completed the study instruments. The results showed that the mean of workaholism, positive perfectionism, and negative perfectionism were respectively 2.60, 4.04, and 3.53. Additionally, multivariate tests showed that the results of post hoc differences for positive perfectionism were in favor of males, subordinates, those with a bachelor’s degree, those with less than 5 years of experience, and those aged less than 30 years. Furthermore, the differences for negative perfectionism were in favor of those with a bachelor’s degree and subordinates. For workaholism, the differences were in favor of subordinates, public sector employees, married persons, and those with a diploma degree. Finally, the results of hierarchical regression analysis found that positive and negative perfectionism and some demographic variables predicted 129% of the variability in workaholism, and the typical hierarchical regression model included positive and negative perfectionism without other demographic variables.

## Introduction

Workaholism is defined as “the tendency towards excessive interest in work and stimulates irrational motivation to invest most of one’s time and effort to work, which hampers other important life activities” (Andreassen, Hetland, & Palesen, 2014, p. 8). The results of some studies have shown that the prevalence of workaholism in European societies ranges from 8 to 10% (Sussman, Lisha & Griffiths, 2011), and only one study so far has examined the prevalence of workaholism around the world. The Norwegian community found that the prevalence of workaholism was 8.3% (Andreassen et al., 2014).

Although workaholism is not a simple percentage, all signs suggest that there will be an increase in the rate of workaholism in the future because of changes in the pattern of work, which has become dependent on smart devices and the internet, and changes in work modes and technology (Andreassen, Griffiths, Sinha, Hetland, & Pallesen, 2016; Quinones, Griffiths, & Kakabadse, 2016). Empirical work has been investigated comparing enthusiastic work and honest work, and there are clear differences in terms of job addiction, competence, and sincerity (Schaufeli, Taris, & van Rhenen, 2008). In fact, efficiency and sincerity in work are linked to increased satisfaction with life, achievement in work, and professional, functional and financial progress. (Anderson, Aziz, & Wuensch, 2016). Workaholism is also associated with psychological stress, incivility, anxiety and depression (Andreassen et al., 2016) and contributes to lower satisfaction with work (Karanika-Murray, Pontes, Griffiths & Biron, Shimazu, Schaufeli, Kamiyama, & Kawakami, 2015). Workaholism leads to psychological burnout, family conflict, and reduced productivity (Griffiths & Karanika-Murray, 2012; Sussman, 2012).

It is a form of behavioral addiction similar to other forms of behavioral addiction, such as addiction to games, gambling, sports and sex (Demetrovics & Griffiths, 2012). As described in Griffiths (2005a), the components of the addiction model consist of seven criteria that contribute to the satisfaction of the addict. These criteria are 1. Importance, i.e., “that work is the most important activity in their lives”; 2. Mood modification, i.e., “any work that is used to relieve emotional stress or to generate thrills and cheerfulness”; 3. Endurance, i.e., “the need to work longer and more intensely to get a cheerful mood”; 4. Withdrawal, i.e., “any sense of emotional stress, suffering and emotional turmoil as soon as the work is interrupted”; 5. Conflict, i.e., “the existence of conflicts and problems with family members and individuals who contradict their lifestyle”; 6. Relapse, i.e., “any return to the old patterns of work after a period of adjustment and drop-out”; and 7. Problems, i.e., “that addiction to work affects health, personal relationships and hobbies and thus causes problems” (Andreassen, Griffiths, Hetland, & Pallesen, 2012; Brown, 1993; Griffiths, 2005b).

Empirically, the measures of workaholism have failed to show factors of workaholism (Andreassen, et al., 2014; Burke, Richardsen & Martinussen, 2002; Kanai, Wakabayashi & Fling, 1996). Some studies have shown that the factor of fun in work was not directly related to work addiction (Schaufeli, Shimazu & Taris, 2009), which suggests that the core factor of workaholism is the compulsive impulse related to the need for action. Many studies returned to the point of view that there truly was a scientific concept called workaholism. This concept is known to meet a number of scientific criteria (Wojdylo, Baumann, Buczny, Owens, and Kuhl, 2013), and these seven dimensions were adopted as the dimensions of this study (American Psychiatric Association (APA), 2013; World Health Organization WHO, 1992).

Although the concept of workaholism and its names have varied from workaholism (Spence & Robbins, 1992) to work addiction (Robinson, 2013), most researchers did not know or measure the concept of addiction in light of what was mentioned above about the seven dimensions. Previous measures such as a battery about workaholism or work addiction considered addiction to be a behavioral trend, obsessive-compulsive disorder or a type A personality (Andreassen, et al., 2012; Schaufeli, et al., 2009). Recently, Bergen and Anderson (Andreassen et al., 2012) developed a measure of workaholism based on Brown’s (1993) measure of behavioral addictions and Griffiths (2005b) description of the components of common addiction models. All questions on the Berger scale were answered on a five-point Likert scale ranging from never (1) to always (5) by asking how much during the past year the person had symptoms of workaholism. If four questions out of seven were scored a 4 or 5, it indicated that the person suffered from work addiction (Lemmens, Valkenburg, & Peter, 2009). This cut-off was adopted in accordance with previous disciplines in behavioral addiction and a number of specialized approaches in psychiatry, such as the fifth Diagnostic and Statistical Manual (APA, 2013; WHO, 1992).

In addition, the cut-off classified the clients into many categories to distinguish the personnel in terms of the ability to respect working hours, leadership status, and attention to the objective public health elements of the scale. This cut-off also distinguishes addicts and non-addicts in light of the number of hours work and levels of exhaustion, and based on the fact that workaholism is more associated with exhaustion and leads to a sense of psychological burnout compared to non-workaholism, all the items of this measure are in the positive direction (Andreassen et al., 2012; Molino, 2013).

On the other hand, Molino (2013) found another study that supported the results of the factor construction of the scale. The results of the validity of the construction confirmed the relation between the scale and job satisfaction, family conflict, workload, cognitive abilities, emotional disorders, and neurotic perfectionism.

Regarding the impact of work addiction on health and psychological issues, many studies have revealed a positive relationship between work addiction and family conflict (Andreassen, Hetland, & Pallesen, 2013; Bonebright, Clay, & Ankenmann, 2000; Russo & Waters, 2006) and between work addiction and psychological stress that lead to psychological and physical symptoms (Andreassen, Hetland, Molde, & Polesen, 2011; Kubota, Shimazu, Kawakami, Takahashi, & Nakata, 2010). For the five personality factors, the results also showed a positive relationship with neuroticism, such that as an individual becomes more nervous and sensitive, more aware, more organized and efficient, workaholism affects his relations with friends and professional colleagues. One study revealed a positive relationship between workaholism and openness (Burke, Matthiesen, & Pallesen, 2006), and other studies have shown that workaholism appears to be unrelated to gender, but it is related to age, especially for young people and adults who spend more time on work (Taris, van Beek, & Schaufeli, 2012).

To date, the percentage of estimates of workaholism varies greatly depending on the sample and tools used, and studies conducted on students found that the rate of workaholism was estimated at 14% (MacLaren, & Best, 2010) when measured by another questionnaire and 18% with subjective estimates (Cook, 1987). In a simple survey of 219 adults by Spence & Robbins (1992), the rate of workaholism was 8% for males and 13% for females. Another study of 192 doctors and psychiatrists found that the proportion of workaholism was 23% (Doerfler, & Kammer, 1986). In another study, the workaholism rate was 21% in 962 Japanese working males (Kanai et al., 1996). In Canada, the results of a study of 519 graduates of the media college showed that the percentage of work addiction was 13% (Burke, 1999). Thus, we can say that the rate of workaholism around the world, in general, ranged from 5% to 10% and was sometimes as high as 25% (Sussman et al., 2011).

Anderson and his colleagues (2014) conducted a study on the rate of work addiction with a survey of a national sample of Norwegian staff. The sample included 1,124 employees. The results showed that the rate of workaholism was 8.3%, and their work negatively correlated with age and was positively related to neuroticism.

In another study with a Brazilian sample of 392 managers, the results showed that workaholism predicted satisfaction with life and that workaholism was a psychological phenomenon that affected the health and well-being of managers and bosses (Pinheiro & Carlotto, 2018).

Liang and Chu (2009) studied the personality traits that cause workaholism in a sample of Chinese employees. The results showed that obsessive-compulsive disorder and a tendency towards achievement, perfectionism and awareness were the main factors and underlying causes of workaholism.

In another study entitled, “Irrational thoughts as a psychological motivation for work addiction from the perspective of health psychologists”, with a sample of eight employees from the educational health sector in Colombia, irrational thoughts, perfectionism, pleading, guilt, fear of losing work, generalized anxiety, the specific characteristics of the institution, the nature of the tasks, the work and the responsibilities were all factors that ultimately led to workaholism (Porras, Velásquez, & Parra, 2018).

In another study entitled, “Addiction to work vs. family conflict among university academics in Norway”, with a sample of 2,186 academics and 2551 administrative staff, the results indicated that the workaholism among academics was higher than in non-academic staff. They also had more work requirements and family conflict. Moreover, workaholism was positively related to family conflict and mediated the relationship between overtime work and family conflict (Torp, Lysfjord, & Midje, 2018).

Girardi, Falco, De Carlo, Dal Corso, and Benevene (2018) found that workload was positively correlated with work addiction. In addition, the results showed that both the amount of work and perfectionism were involved in predicting workaholism among managers. The results showed that there was a positive relationship between workaholism and perfectionism. A high degree of self-direction and exaggerated organization mediated the relationship between perfectionism and workaholism (Stoeber, Davis, & Townley, 2013). In Iran, Booket, Dehghan, and Alizadeh (2018) found that perfectionism, openness, and neuroticism explained 43% of the variation in workaholism among teachers at the Faculty of Medical Sciences at the University of Kermanshah.

Based on previous studies, workaholism was negatively associated with receptivity and openness and was not associated with extraversion. Moreover, addiction was not expected to be associated with gender but could be related to age according to the results of some previous studies.

Despite the partial agreement for the main dimensions of workaholism, some studies still have doubts about the real dimensions of workaholism. For example, Spence & Robbins (1992) reported that workaholism was the same as doing high levels of work with a high degree of motivation, activity and motivation towards work compared to low levels of pleasure at work. A study using factor analysis found two types of workaholism: addiction to work of enthusiasm, which is known as individual achievement with high grades on the three dimensions of the scale of workaholism, and addiction to the feeling of enthusiasm for work, which includes high levels of work and motivation but little pleasure at work. The latter kind of work pattern is the same as that seen during an addiction to real work. This multidimensional measure has been adopted for several criteria that determine the degree of workaholism (Andreassen et al., 2014).

The problem investigated in this study is highlighted by the negative impact of workaholism on a number of aspects of life, such as health, personal, family and productivity aspects, and by the relationship between positive perfectionism (PP) and negative perfectionism (NP) and workaholism in light of a number of variables. This study sought the nature of this relationship. More specifically, this study sought to answer the following questions:

1. What is the level of each type of workaholism, PP and NP considering gender, age, qualifications, professional experience, authority, institutional prescription and social status?
2. There are differences in the means of workaholism, PP and NP that can be attributed to the demographic variables.
3. To what extent do PP and NP predict work addiction, and what is the typical multi-hierarchical linear regression equation for predicting work addiction?

This study is the first Arab and Jordanian study, as there have been no Arab studies conducted on this topic to the best of the researcher’s knowledge. All previous foreign studies have been conducted on ethnically diverse and non-national samples. Previous studies have not relied on the criteria of behavioral addiction or the Diagnostic and Statistical Manual. On the other hand, most of the previous studies have relied on a dimension that was not related to work addiction, i.e., pleasure at work. Moreover, most measures of workaholism were used in studies that have failed to employ cut-off points for addiction categories, although the relationship between workaholism and other variables such as personality traits have been explored in an early research. However, there are still some contradictions between these studies that indicate a need for more research. This study is of importance to research and theory because of its procedures and the accompanying scientific results. It also has practical importance by focusing on the importance of dealing with psychological problems, work pressures and family problems directly associated with workaholism.

The main objective of this study was to explore the level of workaholism along with the level of PP, NP, and the relationships among workaholism and other demographic variables, such as age, gender, social status, qualifications, authority, institutional prescription and professional experience. Additionally, this study aimed to explore how PP, NP and demographic variables predict workaholism.

## Method

### Participants

The study was composed of Jordanian employees in the public and private sectors. There were 686 students who met the requirements of the study, including 281 males (SD = 10.23) and 405 females (SD = 9.51). The mean age ranged from 25 to 62 years and averaged 39.42 years. Public sector employees constituted 75% of the total sample. All respondents admitted that they had a supervisor, a boss or a manager at work. The sample included all individuals working overtime or part-time. Retired individuals and those under the age of 18 were excluded.

### Procedures

The study data were obtained through various social media sites and e-mail. Each person was asked to kindly resubmit the questionnaires to all the individuals involved with him in the group in addition to completing the criteria and sending them back to the originator. The number was circulated to an extra set of individuals by snowball sampling from social networking sites, universities and ministries. In total, approximately 782 letters containing all study tools and demographic variables were sent to groups and staff members. It is worth mentioning that the contents of the questionnaire were sent using the Mourners Recruited system so that the data were automatically downloaded. Consent to participate in the research was obtained by informing the participants that they should send the questionnaire only if the data are correct and that they were not obligated to fill it.

The confidentiality of the information was confirmed, and the results of this study and the information they sent were used for scientific research purposes. There was no way to disclose the identity of a single subject. In accordance with the principles of the Helsinki Declaration, which includes the preservation of the ethics of scientific research, there was no need to obtain formal approval from anybody or scientific research committees, as long as the subjects had the right to refuse and were not authorized to disclose their names or any information identifying them.

### Instruments

The Workaholism Scale (Andreassen et al., 2012). This measure has been used to assess work addiction. The measure consists of seven items measuring the symptoms of workaholism using the criteria of addiction: importance, endurance, mood modification, relapse, withdrawal symptoms, conflict and problems. Participants were asked to remember experiences from the past year and choose whether they experienced the abovementioned symptoms from five options (5-point Likert scale), expressed as sentences and words based on diagnostic criteria for addiction, from 1=never to 5=always. The results showed the validity of the factor structure of this scale, RMSWA= 0.08, CFI=096, TLI=095. For internal consistency analysis, the value was between 80 and 84. These results were based on a sample of 12137 employees, and a score of 4 or higher on four or five of the seven items is typically the cut-off between workaholism and non-workaholism, according to the classification of modern psychiatric diseases. The results using the cut-off showed that the scale could distinguish between those addicted to work and those not addicted to work. Cronbach’s alpha of the internal consistency was .81.

The procedures of the study had several initial steps, including approval by the author and then translating the scale from English to Arabic. The process of translating the scale was started by preparing and translating the English version into an Arabic version, and then the scale was presented to two psychologists who had mastered Arabic and English and two other translation specialists to improve the translation. In a subsequent step, the four versions were compared to each other. The decision was made to paraphrase some items, modify some, and change some words to more precise and specialized options. The Arabic version was then translated back to English and compared to the original version to verify conformity in the meaning and translation. Finally, to make the scale suitable for the Arabic language, in general, and for the Jordanian context in particular, the scale was presented to 10 psychologists, and all of them confirmed that the measure was appropriate in its current form for the Jordanian environment. The correlation coefficients were found to be between .57 and .84. Cronbach’s alpha was .91 for the entire scale and was between .85 and .92 for the subscales. The test-retest reliability with a four-week interval was between .86 and .94.

PP and NP scale (Terry-Short, Owens, Slade, & Dewey, 1995). This scale consists of 40 items divided into the PP and NP factors, each with 20 items. The answers ranged from 1 = strongly disagree to 5 = strongly agree. The scale was separately corrected for each factor, and a high score indicates a high degree of PP or NP. Many studies around the world and in different cultures have shown that the scale has acceptable psychometric properties. The consistency coefficient of internal consistency was 0.88 and 0.92 for PP and NP, respectively.

For the purposes of this study, Aldahadha (2018) standardized this measure in Jordanian society. The results of the exploratory factor analysis showed that the scales had two dimensions with 15 positive and 15 negative points. The results showed that PP, depression and anxiety had correlations of 0.88 and 0.91, respectively, while perfectionism was positively associated with depression and anxiety, with correlations of 0.91 and 0.86, respectively. For the stability of the scale, the results for internal consistency and test-retest reliability showed that the stability coefficient ranged from 0.86 to 0.92 for both PP and NP. Therefore, after correcting the scale for both PP and NP, scores from 1-1.66 indicated no perfectionism, those from 1.67 - 3.33 indicated medium perfectionism, and those from 3.34-5 indicated high perfectionism.

### Statistical analysis

The data were analyzed with SPSS version 24. Reliability and Pearson’s correlations were analyzed, as were descriptive statistics including ratios, means and standard deviations. Univariate variance analysis and hierarchical regression analysis were performed. The independent demographic variables were gender, age, professional experience, institutional prescription, qualification, social status and authority, and the dependent variables were workaholism and PP and NP.

## Results

To answer the first question about the level of workaholism, the means and standard deviations of employee PP and NP grouped by demographic variables were calculated. The cut-off on the scale was adopted to determine those who suffered from work addiction from those who did not. Answering four questions out of 7 with a 4 or 5 indicated that a person was suffering from work addiction (Lemmens, Valkenburg, & Peter, 2009). This cut-off was adopted from previous research in behavioral addiction and the fifth Diagnostic and Statistical Manual (APA, 2013; WHO, 1992). Thus, the cut-off was 16 + 20 ÷ 2 = 18, which was a mean of 2.57. Therefore, means of 2.57 and above on the total scale indicated addiction to work.

Table 1 shows that the means of the workaholism items were around the mean of the cut-off, with a slight increase or decrease, ranging from 2.46 for leaders and those with a diploma to 2.75 for those with 5 years of professional experience or less. As shown in the same table, in general, the mean responses of the sample on the scale were above the cut-off point of 2.60, indicating slight workaholism in the study sample, in general. For the mean responses for PP and NP, the results were within the high range for the two scales as a whole and for demographic variables. The means ranged from 3.43 to 4.17 at the highest level. The total score of PP was 4.02, which was a very high score. For NP, the mean was 3.53, which was also high.

**Table 1.**
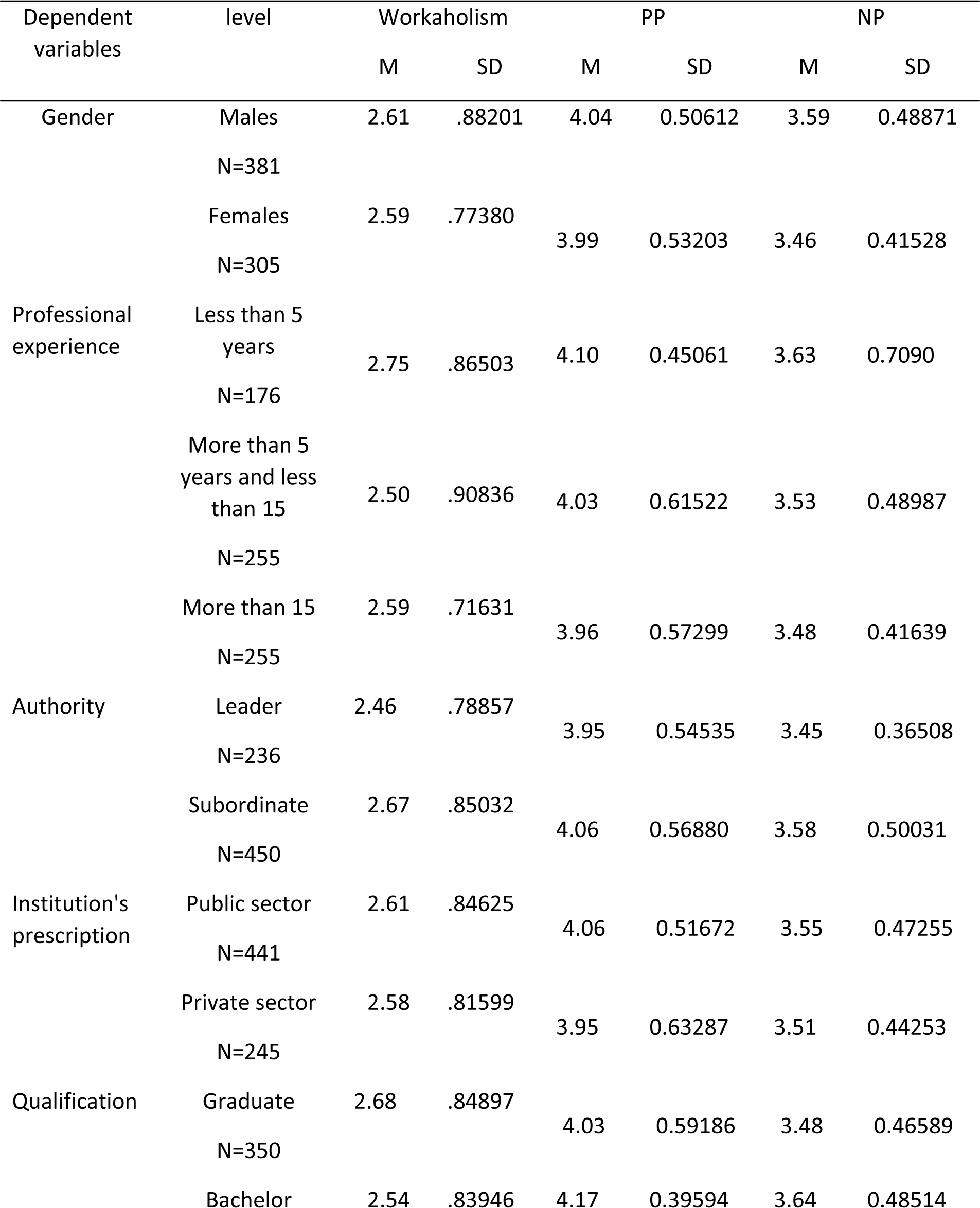
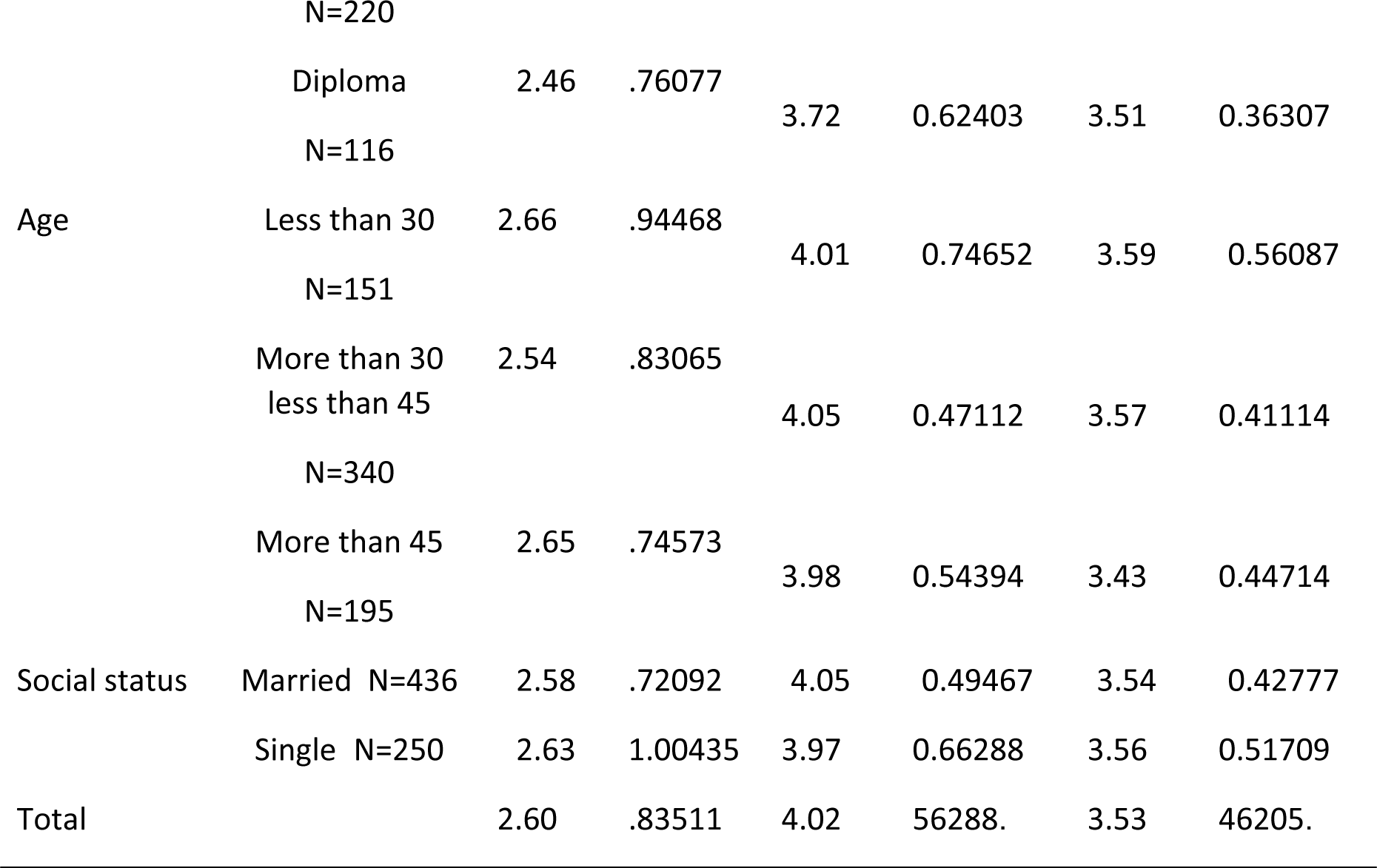
Means and standards deviations due to the study variables

To answer the second question and to check the significance of differences in means statistically, a univariate multivariate test was performed. Table 2 shows the results of multiple variance analysis for the independent and dependent study variables.

**Table 2.**
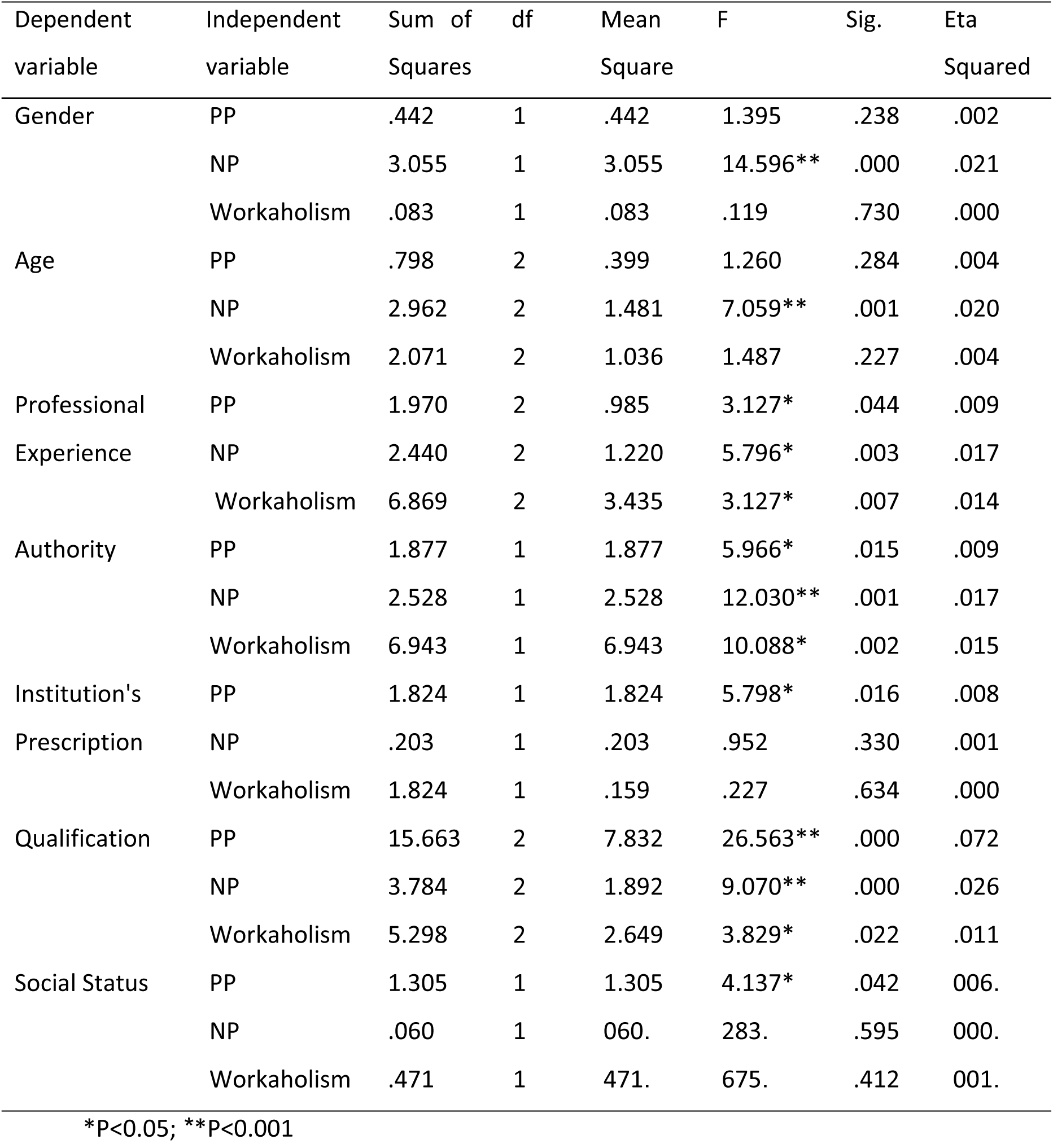
Results of the Univariate Multivariate analysis due to the study variables

| Dependent variable | Independent variable | Sum of Squares | df | Mean Square | F | Sig. | Eta Squared |
| --- | --- | --- | --- | --- | --- | --- | --- |
| Gender | PP | .442 | 1 | .442 | 1.395 | .238 | .002 |
|  | NP | 3.055 | 1 | 3.055 | 14.596** | .000 | .021 |
|  | Workaholism | .083 | 1 | .083 | .119 | .730 | .000 |
| Age | PP | .798 | 2 | .399 | 1.260 | .284 | .004 |
|  | NP | 2.962 | 2 | 1.481 | 7.059** | .001 | .020 |
|  | Workaholism | 2.071 | 2 | 1.036 | 1.487 | .227 | .004 |
| Professional | PP | 1.970 | 2 | .985 | 3.127* | .044 | .009 |
| Experience | NP | 2.440 | 2 | 1.220 | 5.796* | .003 | .017 |
|  | Workaholism | 6.869 | 2 | 3.435 | 3.127* | .007 | .014 |
| Authority | PP | 1.877 | 1 | 1.877 | 5.966* | .015 | .009 |
|  | NP | 2.528 | 1 | 2.528 | 12.030** | .001 | .017 |
|  | Workaholism | 6.943 | 1 | 6.943 | 10.088* | .002 | .015 |
| Institution's | PP | 1.824 | 1 | 1.824 | 5.798* | .016 | .008 |
| Prescription | NP | .203 | 1 | .203 | .952 | .330 | .001 |
|  | Workaholism | 1.824 | 1 | .159 | .227 | .634 | .000 |
| Qualification | PP | 15.663 | 2 | 7.832 | 26.563** | .000 | .072 |
|  | NP | 3.784 | 2 | 1.892 | 9.070** | .000 | .026 |
|  | Workaholism | 5.298 | 2 | 2.649 | 3.829* | .022 | .011 |
| Social Status | PP | 1.305 | 1 | 1.305 | 4.137* | .042 | .006 |
|  | NP | .060 | 1 | .060 | .283 | .595 | .000 |
|  | Workaholism | .471 | 1 | .471 | .675 | .412 | .001 |

\*P<0.05; \*\*P<0.001

The results in table (2) indicate that there are statistically significant differences in the means of workaholism by professional experience, authority, and qualification of 3.127, 10.088, and 2.649, respectively. The results also showed statistically significant differences in the mean of NP by gender, age, professional experience, authority and qualification, with values of 14.596, 7.059, 5.796, 12.030 and 9.070, respectively. The results also showed statistically significant differences in the mean of PP by professional experience, authority, and qualification with 3.127, 5.966, and 26.563, respectively. The eta-squared effect size was found to be small to medium. The effect size can be interpreted as small according to Cohen’s equation (Cohen, 1988). An effect size of 0.01 is small, an effect size of 0.6 is medium, and an effect size of 0.14 is high. To determine the significance of the difference, Tukey’s test was used for multiple comparisons as shown in Table 3.

**Table 3.**
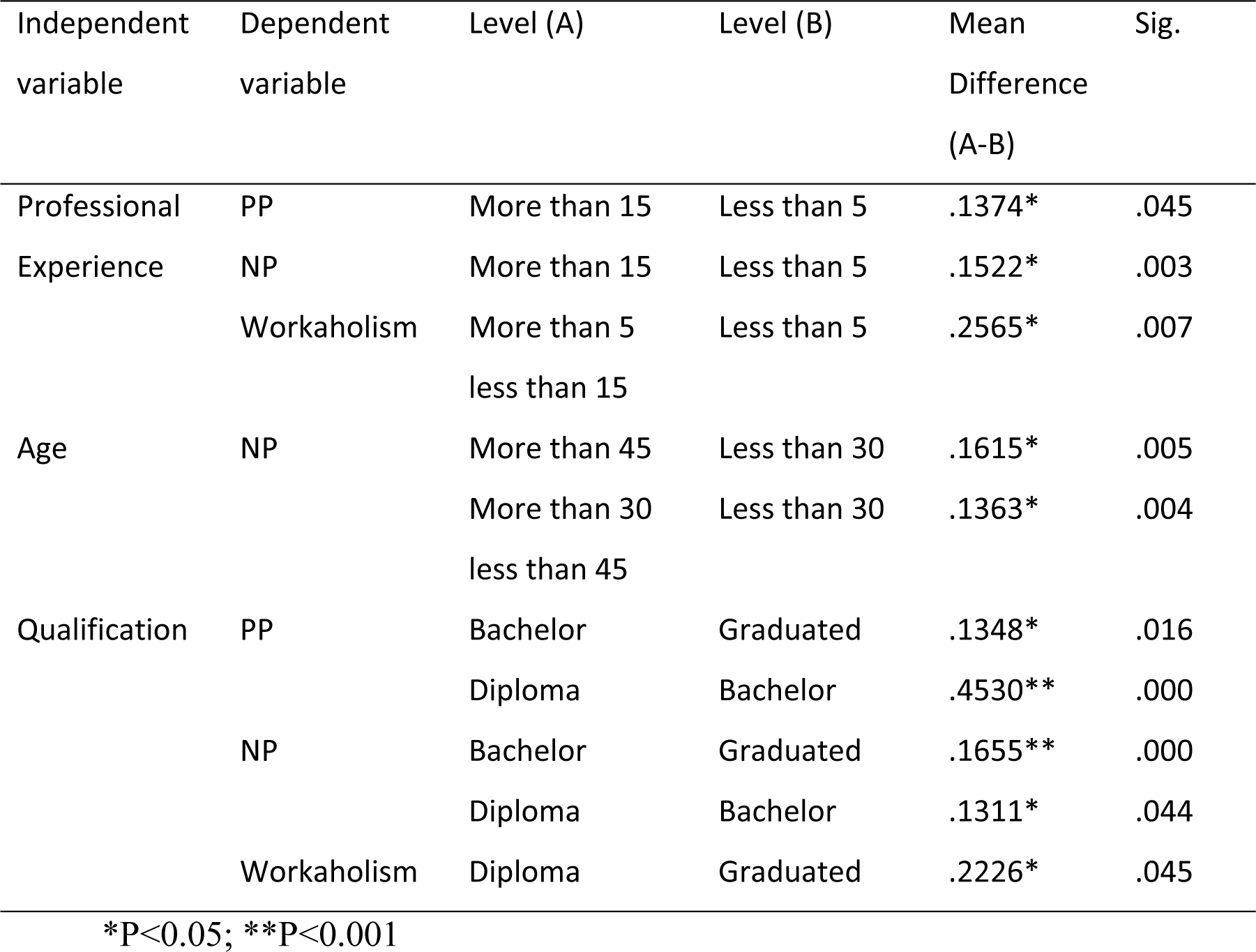
Tokey Post Hoc test of the significant differences between the study variables

The results of Tukey’s test are shown in table 3 and indicate that there are statistically significant differences between the means of PP, NP and workaholism by professional experience, in favor of those with less than 5 years of experience; the means for these factors are 4.10, 3.63, and 2.75, respectively, compared to those with the highest amount of experience. The results of the post hoc comparisons also showed statistically significant differences in the mean of the PP in favor of those aged less than 30 years, with a mean of 3.57, which is higher than that of those who are older. Finally, the results for workaholism favor bachelor’s students for PP (M = 4.17) and NP (M =3.64) and those with a diploma compared with those who graduated (M = 3.51).

In terms of gender, authority, institutional prescription and social status, the results favored males for PP (M = 4.04). The results also favored subordinates for workaholism, M = 3.58, PP (M = 4.0), and NP (M=3.58). On the other hand, the results for workaholism favored the public sector (M = 3.55) and those who were married (M = 3.54).

To answer the third question, linear regression analysis was carried out. PP was added in the first model, and PP and NP were added in the second model. Finally, in the third model, all of the demographic variables, including PP and NP, were included. The results showed that the correlation coefficients among all the demographic variables, the total score on the workaholism scale and the total scores for PP and NP were between 0.54 and 0.89, all of which were significant at p <.05. Table 4 shows the results of the hierarchical regression models.

**Table 4.** Linear multiple regression analysis of independent variables on workaholism.

| Model | R | R Square | Adjusted R Square | Standard Error | R Square Change | F | df1 | df2 | Sig. |
| --- | --- | --- | --- | --- | --- | --- | --- | --- | --- |
| 1 | .234 | .055 | .054 | .81245 | 0.55 | 39.749** | 1 | 684 | .000 |
| 2 | .311 | .097 | .094 | .79473 | 0.42 | 31.844** | 1 | 683 | .000 |
| 3 | .359 | .129 | .117 | .78472 | 0.32 | 3.503** | 7 | 676 | .001 |

Note: Model 1=PP; Model 2=NP and PP; Model 3= PP, NP, gender, age, authority, Institution's Prescription, experience, qualification, and social status; \*\* $P < 0.001$

It is clear from table 4 that there is a strong correlation between PP and workaholism (R = 234) in the first model, and the change of 0.55 is significant, F= 39.749, p <.001. When NP was added in the second model, the correlation coefficient was 0.500, and the change increased very little to 0.42, which was significant, F= 31.844, p> 0.001. In the third model, all demographic variables were added. The change also increased to 0.32. Thus, the R^2^ was 129. This value indicated the variance in workaholism that was explained using the three models together.

Accordingly, table 5, which shows the linear regression equation and function, was statistically extracted after including both PP and NP and excluding all other demographic variables. PP in the first model was t=82.358, which was significant at p <.001; when adding NP in the second model, the value was t=7.400, which was also significant at p<.001. Thus, the hierarchical linear regression equation for predicting workaholism consists of both PP and NP.

**Table 5.**
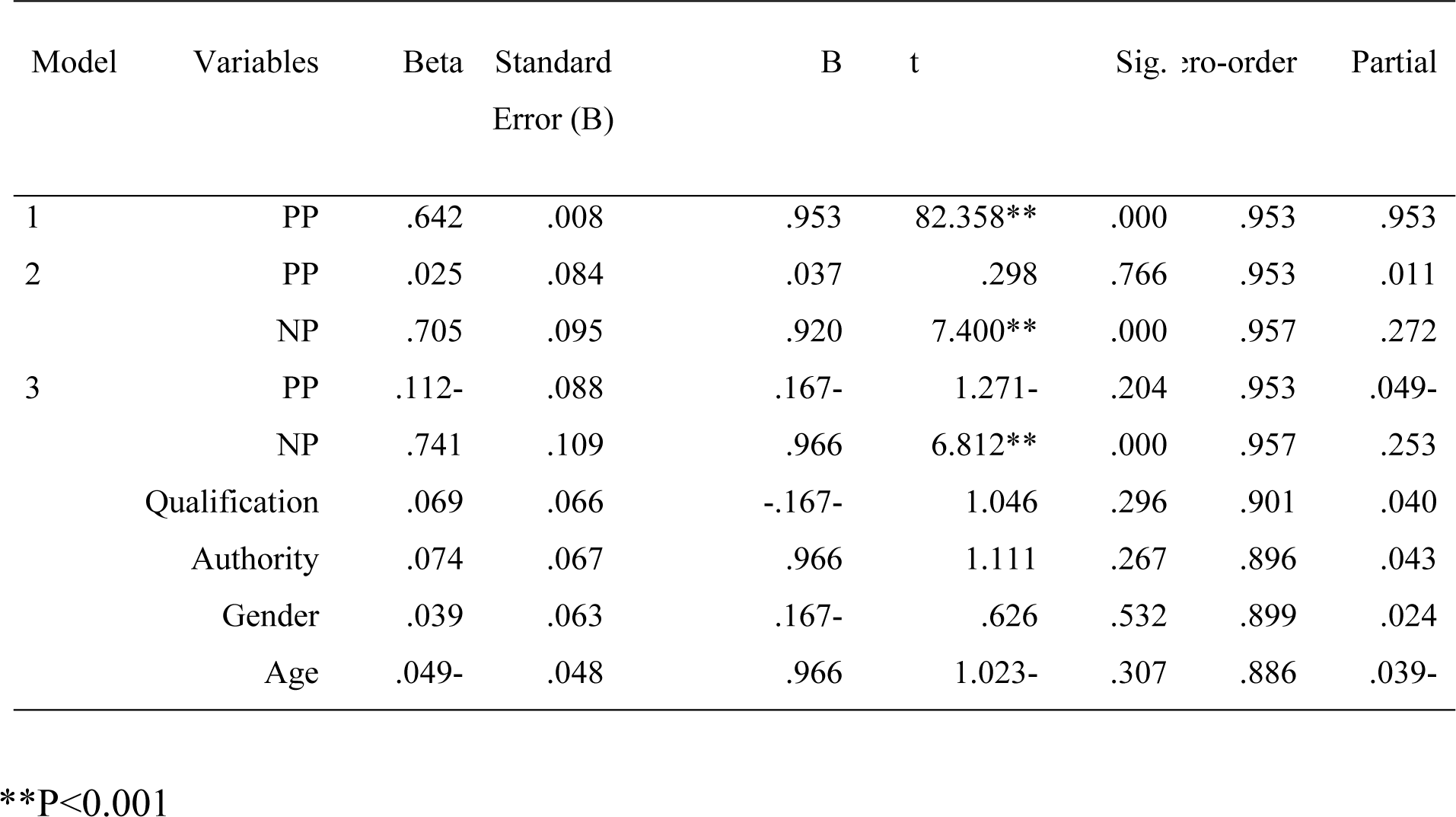
The typical multi-hierarchical linear regression equation for predicting workaholism

The normal distribution of workaholism for the individuals in the study sample was also extracted and is shown in figure 1. The figure shows that workaholism had a normal distribution. Based on the linear model, which also shows that the data were distributed around a straight line, the random distribution condition was achieved.

**Figure 1.**
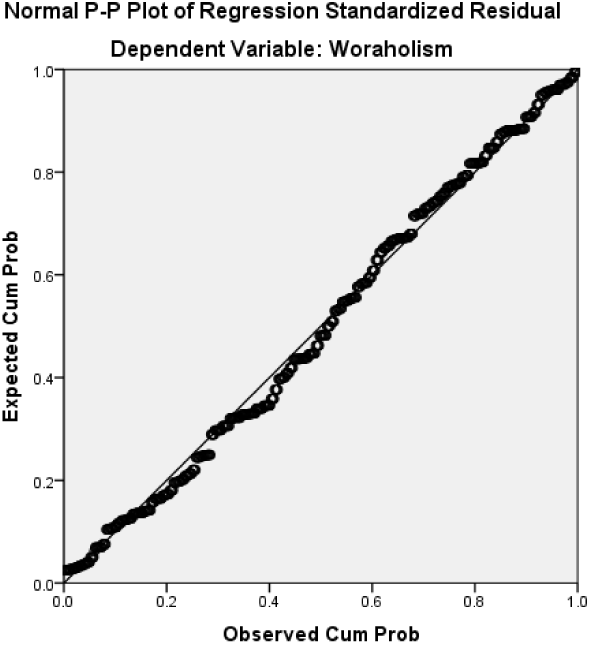
Distribution of the study sample on linear regression of workaholism

## Discussion

This study aimed to explore workaholism, PP, and NP in a sample of Jordanian employees. The sample members recorded a slightly higher mean for workaholism in general. Grouped by demographic variables, the means ranged from 2.46 for both leaders and those with diplomas to the lowest mean of 2.75 for those with 5 years of professional experience or less. The overall mean of 2.60 indicated slight workaholism. For the PP and NP means, the results were within the high range on the two scales overall and when grouped by demographic variables. The means ranged from 3.43 to 4.17, and the mean for the total PP score was 4.02, which was a very high score. For NP, the mean was 3.53, which was also a high value. It seems that both professional leaders and those with academic qualifications with or without a diploma do not suffer from workaholism. This finding is likely due to the fact that the leader has the administrative and accounting powers that allow him or her to judge the achievements of subordinates and those who are lower in the job hierarchy. He or she can feel comfortable and reassured and does not need to do more to convince his boss, unless the leader is at work seeking to please those who are in top management positions or may have a personal style, as indicated by Russo and Waters (2006).

Employees with academic qualifications, such as a diploma, without work addiction had a low average, which explains why this category may not have more precise job centers and suggests that productivity may not be of great value to them. They occupy a functional space only, without need for more creativity that may lead them to work addiction. On the other hand, work addiction was higher for those with 5 years of experience or less, as this group is expected to be a younger age group with a desire to advances. These factors all require more work and motivation.

The results for PP and NP means were high, and this result may be expected from Jordanian employees. This feature has been expressed by some Gulf countries that attract Jordanian employees. The Jordanian worker is characterized by seriousness and perseverance. These are the same traits and symptoms that both PP and NP may indicate, but the difference between them lies in the extremes of NP and its associations with symptoms and cognitive distortions, while PP is associated with productivity and moderation. The results of this study are consistent with studies by Andreasen et al., 2014; Burke, Richardsen & Martinussen, 2002; Kanai, Wakabayashi & Fling, 1996. Most of these studies confirmed that workaholism is a satisfactory concept that has psychological symptoms consistent with the criteria of a modern psychiatric diagnosis. The results of this study are also consistent with the studies of Andreassen, Hetland, Molde, & Palesen, 2011; Kubota, Shimazu, Kawakami, Takahashi, & Nakata, 2010, affirming that addiction has a negative impact on an individual’s health.

The results of the post-hoc comparisons for the multivariate analysis showed that there are significantly different means of PP and NP and workaholism by professional experience in favor of those with less than 5 years of experience compared to those who have the highest amount of experience. The means of those with lower amounts of experience were the highest, and they have a tendency towards excellence compared to those who have the highest amount of experience. In contrast, those with advanced professional experience have been subjected to psychological pressure and burnout, which invites them to refrain from working to the degree classified as workaholism. The optimal degree of work does not fall under the category of workaholism but is instead a moderate degree of work, which is without the pieces that have been previously referred to in this article.

The results also showed statistically significant differences in the mean of PP by age, in favor of those younger than 30 years old, in which the mean was greater than that of those who were older. This result was also consistent with the previous result, as it was obvious that an employee with less than 5 years of experience will be younger than 30 years old. Therefore, the interpretation of this result may not differ greatly from the previous interpretation of those with less than 5 years of experience. For gender, authority, institutional prescription, and social status, the results favored males for PP. This result can be explained by the fact that males endure the greatest responsibility for home, marriage, and financial expenditures in general, and they do more in hopes of higher incomes and a better life. The results favored subordinates for workaholism and measures of PP and NP. This result may be the most interesting, as the subordinates at work progress at all other levels of demographic variables by obtaining the highest means on workaholism and perfectionism, which indicates that the subordinate, compared to the leader, tends to load himself with more work and performs many duties to satisfy the president. On the other hand, his behavior is characterized by PP and NP. Some people do not differentiate between PP and NP, and they instead combine a mixture of behaviors from both scales, causing confusion and ambiguity in who the right person is for different actions.

The results for workaholism were also in favor of public institutions and those who were married, which can be explained by married individuals devoting most of their time to work. Unmarried individuals still have other interests, and the most important interest for the unmarried is to find a future wife or husband. Public institutions sometimes provide additional work, thereby increasing the burden of workaholism. Finally, the results for workaholism favored those with a bachelor’s degree for PP and NP compared to those with a graduate degree and favored those with a diploma compared to those with a graduate degree. This result is also expected and agrees with the results for subordinates, those with less than 5 years of experience and those under the age of 30. All of these groups are within the same age group and can be grouped as trainees. They appear to be the most targeted group for workaholism, PP and NP. The results of this study agreed with Travis, van Beek, and Schaufeli (2012) and Molino (2013), whose results supported the relationship between job satisfaction, family conflict, workload, irrational beliefs, emotional disorders and neurotic perfectionism.

Three models were tested for the results of the hierarchical regression analysis. The PP scale was added in the first model, PP and NP were in the second model, and all demographic variables and PP and NP were included in the third model. The results showed that the correlation coefficients among all demographic variables and the total score of the measure of workaholism and the total scores of the measures of PP and NP were all significant. This result was also expected. The relationship was strong between workaholism and PP and NP. In addition, as interpreted through these two measures, perfectionism is a direct cause and the main catalyst of work addiction for employees (Girardi et al., 2018).

The R^2^ value was 129. This value is the variance in workaholism that is explained by all three models combined. Thus, the regression equation for the linear model was statistically derived using PP and NP with all other demographic variables excluded. Therefore, the interpretation of workaholism is largely explained by PP and NP.

### Conclusion and recommendations

This study has been an important step in uncovering a real problem that has not been discussed previously in Arab societies. A global scale was introduced for the first time in an Arab and Jordanian environment. This study makes important contributions by providing psychologists with new facts about the importance of focusing on workaholism symptoms. However, this study has some limitations and strengths. The sample was not patients and was instead made up of general staff. Therefore, we recommend new studies with samples of patients with mental disorders, and conducting studies on specific groups of staff with obsessive-compulsive disorder. The results of this study are determined by the extent of the validity of the responses of the sample members to the criteria. The diagnosis was not according to the clinical interview. On the other hand, the scale of workaholism was consistent with the criteria of addiction according to the statistical diagnostic guide.

